# Seed2LP: seed inference in metabolic networks for reverse ecology applications

**DOI:** 10.1101/2024.09.26.615309

**Authors:** Chabname Ghassemi Nedjad, Mathieu Bolteau, Lucas Bourneuf, Loïc Paulevé, Clémence Frioux

## Abstract

A challenging problem in microbiology is to determine nutritional requirements of microorganisms and culture them, especially for the microbial dark matter detected solely with culture-independent methods. The latter foster an increasing amount of genomic sequences that can be explored with reverse ecology approaches to raise hypotheses on the corresponding populations. Building upon genome scale metabolic networks (GSMNs) obtained from genome annotations, metabolic models predict contextualised phenotypes using nutrient information. We developed the tool *Seed2LP*, addressing the inverse problem of predicting source nutrients, or *seeds*, from a GSMN and a metabolic objective. The originality of Seed2LP is its hybrid model, combining a scalable and discrete Boolean approximation of metabolic activity, with the numerically accurate flux balance analysis (FBA). Seed inference is highly customisable, with multiple search and solving modes, exploring the search space of external and internal metabolites combinations. Application to a benchmark of 107 curated GSMNs highlights the usefulness of a logic modelling method over a graph-based approach to predict seeds, and the relevance of hybrid solving to satisfy FBA constraints. Focusing on the dependency between metabolism and environment, Seed2LP is a computational support contributing to address the multifactorial challenge of culturing possibly uncultured microorganisms. Seed2LP is available on https://github.com/bioasp/seed2lp.

## Introduction

In the past decades, metagenomic sequencing had a profound impact on microbiology with substantial expansion of known genome collections [Zeng et al., 2022, Nayfach et al., 2020, Nishimura and Yoshizawa, 2022]. While the previously widespread paradigm that only 1% of micro-organisms are culturable tends to be rejected [Martiny, 2019], estimations state that a high proportion of archaea and bacteria remain uncultured across most biomes [Steen et al., 2019]. The prevalence of microbial interactions in natural environments is a strong hypothesis underlying the difficulty to culture many taxa [Yan et al., 2023], and more and more studies pinpoint the role of metabolism in setting up these interactions [Pande and Kost, 2017, Pacheco et al., 2019]. While microbiology is essential to solve the problem of culturability, computational methods taking advantage of the availability of genomes coupled to simulation approaches can provide complementary hypotheses to tackle this issue.

Computational models of metabolism enable the prediction of metabolic activity in organisms, taking into account a description of environmental conditions and additional constraints. In practice, the information related to the biochemical reactions a microbe could catalyse is abstracted in genome-scale metabolic networks (GSMNs), that are in turn used in models for simulation, such as flux balance analysis (FBA) [Orth et al., 2010]. GSMNs can be automatically or semi-automatically built starting from genomes [Gu et al., 2019], although it is unrealistic to expect a high quality GSMN based on a fully automatic process [Karp et al., 2018]: manual curation, expert knowledge and refinement are needed to reach a level of quality sufficient for quantitative simulations of the model [Thiele and Palsson, 2010, Bernstein et al., 2021]. As a result, it is much more difficult to model the metabolism of poorly-studied organisms that are sometimes impossible to experimentally grow in pure culture and defined medium, and for which only a (possibly incomplete) genome is available. For these cases, quantitative models of metabolism are hardly applicable in first intention. Topological analysis of the graph structure of GSMNs and qualitative simulations are then a good trade-off to gain knowledge on possibly incomplete models [Biggs et al., 2015, Bosi et al., 2017], and improve them prior to quantitative simulations [Bernstein et al., 2019].

Using GSMNs to predict the growth environment of a species has previously got attention. Three classes of approaches can be distinguished: structural methods relying on the topology of the graph, reachability methods enabling a qualitative abstraction of metabolic activation, and constraint-based ones ensuring steady state modelling. In the first category, NetSeed [Carr and Borenstein, 2012] calculates the minimal subset of exogenously acquired metabolites, or *seeds* using the decomposition of a directed simple GSMN graph into strongly connected components (SCC). A SCC without an incoming edge is a source component among which a single seed can be identified. An implementation of these concepts in a R package for reverse ecology is presented in Cao et al. [2016], a Python implementation in Hamilton et al. [2017]. Romero and Karp [2001] first introduced a notion of qualitative graph-based activation as a means to predict the set of compounds that can be produced in a GSMN, regardless of their quantity, solving a forward propagation problem, and the backtracking problem identifying precursor compounds to produce essential metabolites. The forward identification of activable pathways and producible metabolites in GSMNs was later included in the network expansion algorithm [Ebenhöh et al., 2004] as a means to decipher the *scope* of a GSMN starting from a set of seeds metabolites, i.e., available nutrients. The scope encompasses all the metabolites that can be reached from the seeds, regardless their stoichiometry in reactions. In line with this Boolean abstraction of metabolic producibility, Handorf et al. [2008] developed a greedy algorithm for the reverse scope problem, i.e., identifying precursors or seeds for a GSMN, by iteratively reducing the set of seed metabolites. A logic programming implementation using Answer-Set Programming of reverse scope has also been proposed by Schaub and Thiele [2009]. As an alternative, Cottret et al. [2008] and Acuña et al. [2012] provide algorithms computing precursors of a GSMN, with the originality of taking into account self-regenerating cycles, thus being less stringent than network expansion and the scope towards the initiation of reachability from seeds. Lastly, constraint-based approaches involve mixed-integer linear programming (MILP) and linear programming (LP) to ensure steady state. The MENTO algorithm [Zarecki et al., 2014] calculates minimal environments for genome-scale metabolic models, ensuring production of biomass, although no implementation is currently available. If a set of exchange reactions is pre-defined in the model, COBRApy [Ebrahim et al., 2013] implements two approaches: solving a LP problem that minimises their corresponding sum of flux, and a MILP formulation that minimises the number of activated exchange reactions, such as in the EMAF approach [Santos et al., 2017]. SASITA [Andrade et al., 2016] also solves a MILP problem identifying sets of precursors which enable target metabolite production from the GSMN’s hypergraph while constraining the system with the steady state assumption.

The complexity of qualitatively identifying minimal sets of seeds in GSMNs, i.e., the inverse scope problem, has been discussed by Nikoloski et al. [2008], Liu et al. [2009] and Damaschke [2015]. The problem is NP-complete even in acyclic directed graphs. As the search for seeds is not limited to highly curated GSMNs and can be a pre-requisite to reach such level of quality, their inference must be suitable to draft metabolic networks, which advocates for the use of topological or qualitative methods. However, the former have multiple limitations: no flexibility in the metabolic objective, simplification of the GSMN as a simple graph, and no consideration of biochemically feasible reactions [Takemoto and Aie, 2017]. Addressing the combinatorics of seed inference qualitatively can tackle the complexity of the problem, but flux analysis and steady state assumption guarantee are further objectives of the modelling.

In this paper, we address the inverse scope problem: identifying sets of seeds in metabolic networks that guarantee the reachability and steady-state production of molecular targets. We provide a hybrid solving method that ensures the seeds enable the activation of an objective reaction under the steady state hypothesis in FBA. We tested the scalability and relevance of our approach on 107 high-quality GSMNs from the BiGG database [King et al., 2016] demonstrating that a qualitative approach based on knowledge representation and reasoning is a good trade-off between computational efficiency and quality of the predictions. We provide a Python library, Seed2LP, that implements the solving using logic programming and permits customising the parameters of the problem to fit the most use-cases in environment prediction from metabolic networks.

## Background

### Metabolic networks

A metabolic network is a collection **R** of reactions *r* of the form

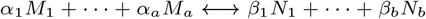

where *M*_1_, …, *M*_*a*_ and *N*_1_, …, *N*_*b*_ are metabolites, and *α*_1_, …, *α*_*a*_ and *β*_1_, …, *β*_*b*_ are stoichiometric coefficients. To each reaction *r* is associated a lower and upper flux bound *ν*_*min,r*_, *ν*_*max,r*_ ∈ ℝ. Whenever one of the bounds is null, the reaction is said irreversible.

#### Flux Balance Analysis

Flux Balance Analysis (FBA) is a widely used mathematical model that aims at predicting flux distributions under a quasi steady state assumption for intracellular metabolites [Orth et al., 2010]. Taking into account additional thermodynamic constraints bounding the flux values for each reaction, FBA solves a linear optimisation problem that maximises (or minimises) the flux in an objective reaction *r*_*T*_ ∈ **R**, usually the biomass reaction that models growth.

The FBA formulation relies on the *stoichiometric matrix* of a metabolic network, which is a matrix 𝒳 ∈ ℝ^|**M**|×|**R**|^, where rows are the metabolites **M**, and columns the reactions **R**. Given a metabolite *m*_*i*_ ∈ **M** and a reaction *r*_*j*_ ∈ **R**, the matrix stoichiometric coefficient 𝒳_*i,j*_ indicates the net gain in metabolite *m*_*i*_ whenever applying reaction *r*_*j*_ positively. To ease notations, we assume hereafter that all metabolites are internal, i.e., each of them is produced and consumed by at least one reaction. The FBA problem is then formulated as follows, with *ν* ∈ ℝ^|**R**|^ a vector of the reaction fluxes:

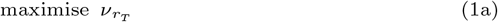

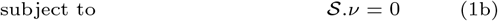

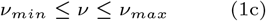

A widely used exchange format for metabolic networks is the Systems Biology Markup Language (SBML) [Hucka et al., 2018] that is used by most metabolic modelling tools. Among the latter, COBRApy [Ebrahim et al., 2013] solves the FBA problem (1) by taking into account data-driven physicochemical and biological constraints. Those can be represented in the form of reactions that hold a special status within the metabolic network as they represent boundaries of the modelled system. By construction, they are unbalanced pseudo reactions, with the objective to add or remove a metabolite for modelling purposes. An exchange reaction *r*_*ex*_ is a reversible reaction with no product, adding and removing an extracellular metabolite *m*_*e*_ ∈ **M**. It is usually of the form *m*_*e*_ ⟷ ∅, with a flux value 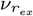 having a negative lower bound and a positive upper bound. The extracellular medium composition that contains nutrients necessary for cell growth will typically be represented as a collection of exchange reactions. A demand reaction *r*_*dm*_ is an irreversible reaction that consumes an intracellular or cytosolic metabolite *m*_*c*_ ∈ **M**: *m*_*c*_ −→ ∅, its flux 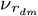 having a null lower bound and a positive upper bound. Lastly, a sink *r*_*sk*_ is a reversible reaction without products, adding and removing a cytosolic metabolite *m*_*c*_ ∈ **M**: *m*_*c*_ ⟷ ∅ with a flux value 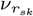 having a negative lower bound and a positive upper bound.

#### Network expansion

The network expansion (NE) algorithm [Ebenhöh et al., 2004] makes a Boolean approximation of metabolic activity from compounds available in the environment. Starting from a subset of metabolites **S** ⊆ **M**, called *seeds*, the algorithm determines a metabolic potential, called *scope*, by inferring the activation of a reaction from the availability of all its reactants, either from seeds or products of other previously activated reactions.

The scope of a metabolic network **N** can be computed iteratively from a set of seeds **S** as described in Eq. (2) until it reaches a fixed point. To ease notations, we assume that reactions are irreversible, and write *reactants*(*r*) and *products*(*r*) the reactants and products of a reaction *r*.

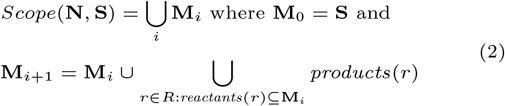

#### Answer-Set Programming

Answer-Set Programming (ASP) is a logic programming paradigm suitable for knowledge representation and reasoning [Baral, 2003]. It enables solving combinatorial problems on Boolean variables subject to logical constraints. An ASP programme consists in a set of atoms, forming the knowledge base, and rules enabling to express deductions, choices, and constraints. Clingo is an ASP solver [Gebser et al., 2019] that was previously used in the context of metabolic modelling applications [Frioux et al., 2018, Thuillier et al., 2021] and can be hybridised to verify linear constraints with clingo-lpx^1^, enabling implementing Boolean constraints such as those from network expansion [Schaub and Thiele, 2009], and checking the satisfiability of linear constraints from FBA, as previously demonstrated in Frioux et al. [2019]. This programming approach permits the implementation of optimisations such as subset minimal or minimal sets of solutions, and offers the possibility of implementing various solving heuristics for the exploration of the solution space by adding constraints and going back and forth with the solver.

## Contributing method

### Model-network reconciliation

The representation of a metabolic model in the SBML format can show inconsistencies between the description of the mathematical model and the structure of the network. As our method relies on both descriptions, it necessitates a normalisation step to reconcile them, particularly in terms of reactions boundaries or reversibility.

A first step is to split reversible reactions into two irreversible ones, thereby ensuring a consistent definition of reactants and products in the scope definition. E.g., a reaction *r*: *A* ↔ *B* with 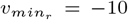 and 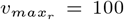 is split into reactions *r*^′^: *A* → *B* [0,100] and 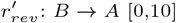. Reactions having a negative lower bound flux and a null upper bound are considered as written backwards because they consumes the metabolites set as products: e.g., *A* → *B* [-100,0] becomes *B* → *A* [0,100]. Reactions having both null upper and lower bounds are deleted. Finally, as existing exchange and sink reactions will artificially create seeds during the inference, it is possible to block the import of the corresponding metabolites: e.g., **A** ↔ ∅ [-10,1000] becoming **A** → ∅ [0,1000], or optionally to maintain a second reaction ∅ → **A** [0,10]. Figure 1a illustrates the above transformations on a simple metabolic network.

**Fig. 1.**
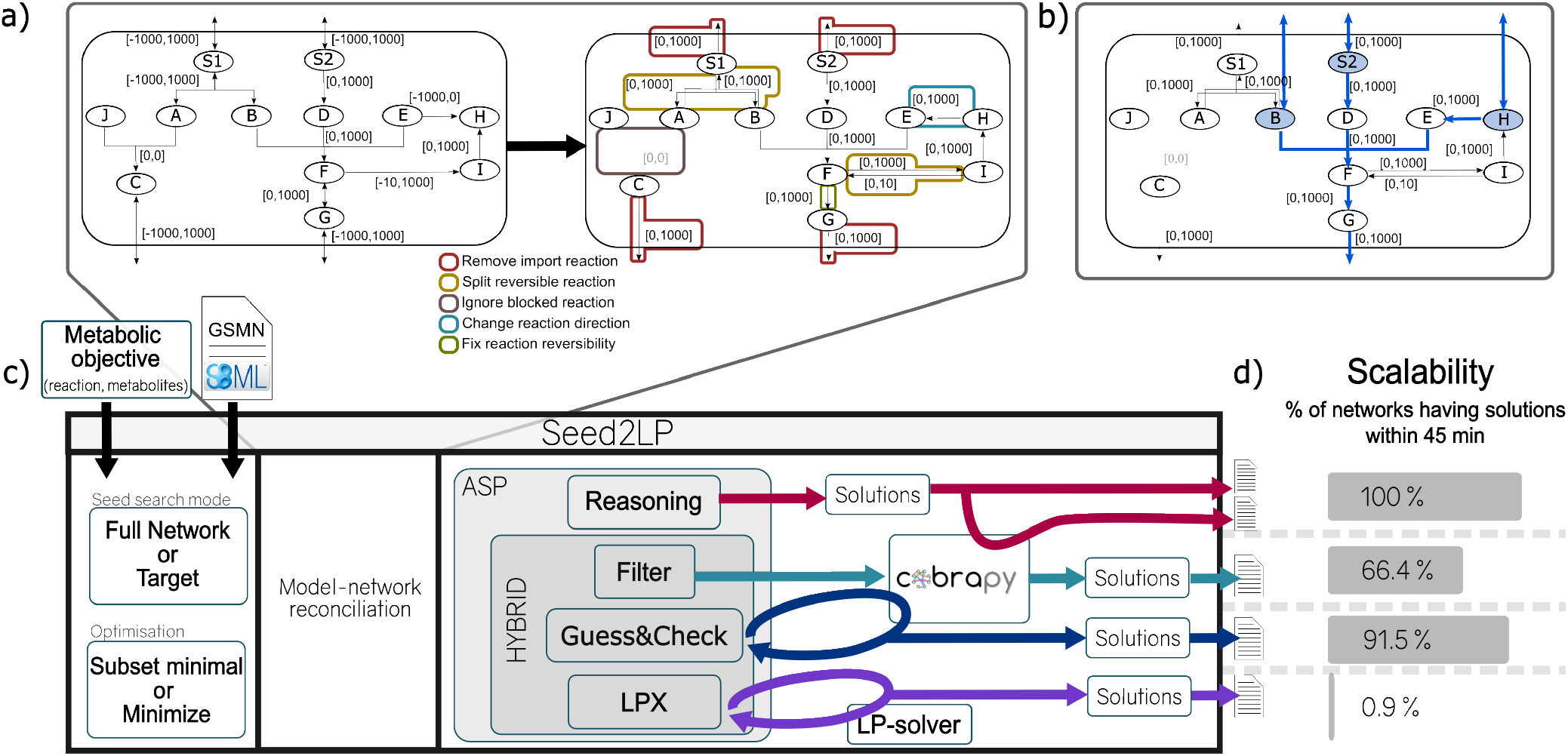
Overview of Seed2LP. **a)** Network normalisation transformation. The first network is the original example, and the second is the normalised network with all modifications highlighted: i) reversible reaction are divided into 2 reactions, ii) reactions with negative boundaries are reversed, iii) import directionality of exchange reactions is blocked but export is maintained, and iv) reactions with boundaries to 0 are deleted. **b)** One seed solution for the previous GSMN with *{B, S*2, *H}*. During COBRApy validation, import reaction is reactivated for S2, sink reactions are created for B and H. Activated reactions are highlighted in blue. **c)** Illustration of Seed2LP. Inputs are a GSMN (SBML) and a metabolic objective. The two main modes are *Full Network* (all metabolites must be reachable) or *Target* (a set of metabolites must be reached). Optimisations for the size of seed sets are subset minimal or minimise. Model-network reconciliation precedes seed inference, for which several solving modes are available. All methods use ASP; reasoning does not include FBA but it can be checked *a posteriori*. The three other methods are hybrid: *Hybrid-filter* returns the n first reasoning solutions validating FBA with COBRApy. *Hybrid-GC* checks flux during solving and eliminates supersets of solutions from the future ones. *Hybrid-lpx* uses a dedicated hybrid solver. **d)** Seed2LP can be run with a time limit, highlighting the scalability of each solving mode, here on the 107 GSMNs from the BiGG database.

In the following, we fix a *normalised* metabolic network **N** over metabolites **M** and reactions **R** and bounds *ν*_*min*_ and *ν*_*max*_.

### Seed search

Seeds are a subset **S** of the network’s metabolites **M** that ensure the satisfiability of a metabolic objective. Such an objective can be defined as a set of metabolites or as a reaction *T* = {(*m*_*obj*_ ⊆ **M**) ∪ (*r*_*obj*_ ∈ **R**)}. The constraints for satisfiability of the objective vary depending on the formalism used for modelling: NE (Eq. (3d)) and/or FBA (Eq. (3a)-(3c)):

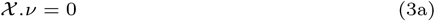

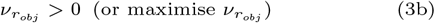

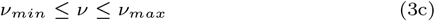

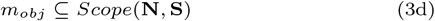

In practice, if metabolites are set as targets, a special case where *T* = {**M** ∪ **R**} is referred to as “full network” mode, leading the reachability of the entire set of metabolites to be the metabolic objective. Alternatively, while setting a metabolic reaction *r*_*obj*_ as an objective is straightforward in FBA, it leads to *reactants*(*r*_*obj*_) to be set as targets in the NE formalism. After seed inference, modelling the presence of seeds in the GSMN implies adding an exchange (respectively a sink) reaction for each seed if the corresponding metabolite is extracellular (respectively cytosolic) in the case constraint-based simulations are performed. Seed inference can rely on the NE formalism or hybrid NE-FBA formalism. We detail below their characteristics.

#### Reasoning-based seed inference

Reasoning-based seed inference aims at identifying sets of seeds **S** satisfying Eq. (3d) for the normalised GSMN **N** under the definition of the scope in Eq. (2). In the case where *T* = {**M** ∪ **R**^′^}, external metabolites of the GSMN are identified as reactants of reactions that are never produced, or solely through transport reactions, and assigned to seeds to initiate scope calculation; remaining seeds being selected from the rest of the metabolites absent from this initial scope. For other cases (*T* = {*m*_*obj*_ ∈ **M**}), the problem is solved in ASP by describing a reverse scope problem identifying a set of putative seed candidates starting from the reactants of reactions producing targets until it reaches a fixed point: seeds are subsets of those:

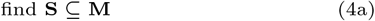

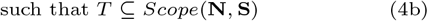

The reasoning mode optimises the size of seed sets in order to reduce the solution space: minimal-size or subset-minimal size seed sets are computed by the solver. The second guarantees that no subset of a inferred seed set can be a solution itself. Given the redundancy of metabolism and the combinatorics of the inverse scope problem, multiple solutions of the same size are likely to exist and can be enumerated.

Seed inference with NE does not ensure the satisfiability of steady state constraints in Eq. (1b), and may permit the accumulation of metabolites (Fig. 1A). Eq. (1b) can be qualitatively approximated by an additional, optional, constraint on the seed search, ensuring that all metabolites in the scope have to be consumed by at least one reaction whose reactants are in the scope (*no-accumulation* mode):

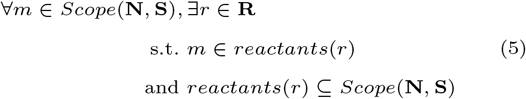

#### Hybrid NE-FBA seed inference

Three flavours of hybrid NE-FBA solving, satisfying constraints from Eq. (3), are proposed. The first one, *Hybrid-lpx* uses Clingo-lpx, combining Clingo to a simplex solver for checking satisfiability of linear equations. Reasoning-based seeds are generated by the ASP solver and their ability to sustain LP constraints are checked within the simplex part of the solver.

The second flavour is *Hybrid-filter* that consists in computing a number of solutions with the reasoning-based search and filtering them *a posteriori* using COBRApy [Ebrahim et al., 2013] (see section External validation with COBRApy): only solutions satisfying Eq. (3) are provided to the user. The set of solutions obtained in *Hybrid-filter* is a subset of the results obtained with the reasoning method.

Finally, a guess-and-check approach *Hybrid-GC* checks the satisfiability of LP constraints in solutions with COBRApy, directly in the process of solving, and adds new constraints to the solver in order to learn from the outcome of the check. Similarly to *Hybrid-lpx*, all solutions outputted by the solver therefore guarantee the steady state assumption and a positive flux in the objective reaction (3). A widened exploration of the solution space that we refer to as *Hybrid-GC*_*div*_ can be obtained by tuning heuristics of the solver to drive it towards diverse solutions.

#### External validation with COBRApy

Whether from *a posteriori* seed inference with the reasoning mode or *Hybrid-filter*, or during solving in *Hybrid-GC*, COBRApy [Ebrahim et al., 2013] solves the FBA problem and assesses whether the seed sets enable a positive flux in the biomass reaction. First, the exchanged reaction are identified and the boundaries are changed to remove the existing imports if any, so that the inferred seeds will be the unique inputs to the model. For all seeds in a solution, the programme will either reopen the import flux bound of its corresponding exchange reaction if any, or create a sink reaction otherwise. The flux in the objective function is then maximised and satisfiability of Eq. (3) is ensured if a positive flux is obtained.

### Implementation

The above methods are implemented in a Python package called Seed2LP. The input is a GSMN in SBML format which will be normalized (see section Model-network reconciliation) prior to seed inference. Existing exchange reactions that will act as seeds can be kept or discarded (default behaviour).

Seed2LP can be run with three different seed searching modes: *FBA mode* which infers seeds with a naive, random selection, by ensuring only the flux into the objective reaction, *Full Network* mode which ensures that all metabolites are in the scope, and the *Target* mode that focuses on a subset of metabolites. The *Target* and *Full Network* modes can be run with five different seed inference methods: *Reasoning, Hybrid-lpx, Hybrid-filter, Hybrid-GC* or *Hybrid-GC*_*div*_ (See section Hybrid NE-FBA seed inference). An objective reaction has to be provided for all modes involving the NE-FBA formalisms. If not set otherwise, default metabolic targets will be the reactants of the objective reaction described in the SBML input file. Figure 1 illustrates the impact of reconciling the network structure and its associated mathematical model in the SBML input, highlights the flexibility of seed search and solving modes of Seed2LP, and illustrating one solution on a toy example.

Additional constraints for seed inference can be provided to the tool: a set of forbidden seeds that can never occur in solutions, a set of existing seeds that will be completed, or a set of possible seeds among which the search will be performed. In the default behaviour, targets are excluded from the seeds to prevent trivial solutions.

Seed2LP provides (subset-)minimal size solution seeds in a json file. It implements FBA post-validation (Eqs. 3a-3c), which is particularly useful for reasoning-based seed inference that does not guarantee steady state. This validation can be used as a standalone step and is complemented by the possibility to assess the size of the NE scope from a given set of seed using MeneTools [Belcour et al., 2020, Aite et al., 2018], thereby verifying that Eq. (3d) holds.

## Results

### A flexible tool for seed inference in metabolic networks

Seed2LP is Python package that infers seed metabolites, i.e., input nutrients, that would enable a metabolic objective to be sustained in a metabolic network according to a simulation paradigm. The core of Seed2LP functioning is machine-reasoning with ASP, and a simplification of metabolic reachability with the network expansion (NE) algorithm. Nonetheless, it implements various hybrid FBA-NE solving modes in order to ensure the resulting models with the inferred seeds satisfy FBA constraints (Fig. 1).

Seed inference was performed on 107 curated GSMNs from the BiGG database [Norsigian et al., 2019] in order to validate the relevance of Seed2LP and results were compared to other approaches (see Methods). Results are described below.

### The Boolean approximation metabolic modelling of Seed2LP outperforms graph-based analysis for seed inference

NetSeed [Carr and Borenstein, 2012] uses the graph structure of the GSMN (see Methods) to identify non-producible compounds called seeds. As it works at the genome scale and does not permit inferring seeds for a subset of target metabolites, it was compared to Seed2LP in the *Full Network* setting. In order to compare the graph-based approach of NetSeed to the relevance of a qualitative model of metabolic producibility (NE), we ran Seed2LP with the reasoning mode and assessed the quality of seed prediction by verifying the possibility to hold a positive flux in the biomass reaction in FBA. A maximum of 1,000 seed solutions were computed for each GSMN by each tool with 45 min timeout for Seed2LP, and with no timeout for NetSeed, which never lasted more than a few minutes for computation.

Results are depicted in Figure 2. NetSeed finds at least a solution set for all 107 GSMNs, while Seed2LP times out for 2 of them. In average, 848 ± 349 and 889 ± 282 solutions could be computed by NetSeed and Seed2LP respectively out of the 1,000 upper bound set up in the benchmark. When applying FBA to solutions computed by NetSeed and Seed2LP, we observed a consistency at the GSMN level: either all solutions for a given GSMN held a positive flux in the biomass reaction or none did. Over the 105 GSMNs instances processed with Seed2LP, 104 have seeds which validated FBA, whereas only 11 did with NetSeed (Fig. 2a).

**Fig. 2.**
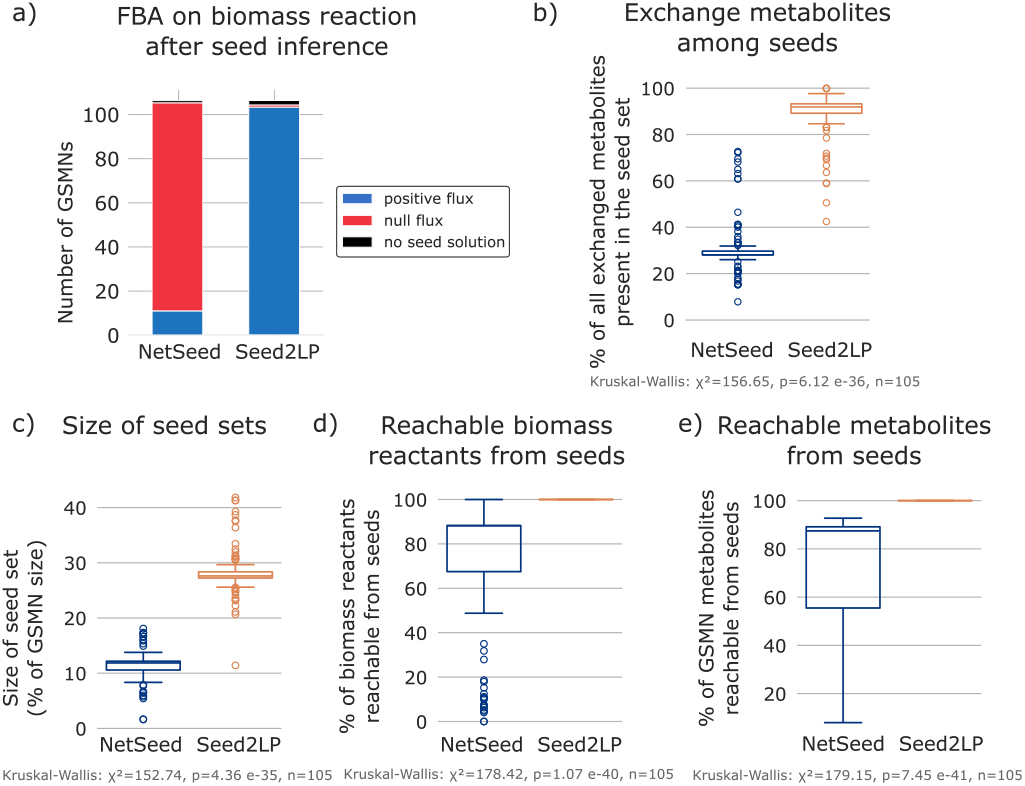
Seed inference with Seed2LP and NetSeed. a) Number of networks having all solutions validating FBA (blue) or not (red) or without any solution (black). b) Proportion of all original exchange metabolites among the inferred seeds. c) Size of seed sets described as the proportion of the total number of metabolites per GSMN. d) Proportion (%) of reachable biomass reactants from the seeds out of the complete set of biomass reactants. e) Percentage of all GSMN metabolites reachable from the seeds. b), c), d), e): Distribution of the average proportions computed by GSMN if several solutions were obtained for a given GSMN. Only results from the 105 GSMNs for which both NetSeed and Seed2LP find solutions are plotted.

We then looked at the coverage of exchange metabolites, i.e., the medium used for simulation as described in the original BiGG GSMNs, by seeds (Fig. 2b). Seed2LP includes a higher proportion of such metabolites than NetSeed (average of averages by GSMN of 89.4% vs 30.6% respectively). Sets of seeds computed by Seed2LP are also larger than those of NetSeed (average of averages by GSMN 28.2% vs 11.5% of the total number of metabolites in the GSMN), which is expected in a context where the entirety of metabolites are expected to be reached (Fig. 2c).

We then surveyed the ability for seeds computed by NetSeed to reach metabolites of the GSMN or biomass precursors using the NE algorithm. By design, all metabolites, therefore including the biomass reactants, are reachable from the seeds computed by Seed2LP. Only 69.4% and 68.6% in average of biomass reactants and of all metabolites respectively were reachable from the seeds proposed by NetSeed (Fig. 2d, e). This indicates that while NetSeed infers seeds from the global GSMN, the purely graph-based solving, and its associated simplifications (e.g. GSMN transformed into a simple graph: a reaction *A* + *B* → *C* is transformed into two reactions *A* → *C* and *B* → *C*) do not enable the reachability of all nodes. On the contrary, the modelling approach of NE as used in Seed2LP ensures this reachability and, despite being qualitative as well, is more consistent with FBA numerical constraints.

We additionally tested the impact of the network normalisation step on NetSeed, performed as pre-treatment by Seed2LP. It appeared that seeds passing FBA validations were identifcal regardless of the normalisation.

Lastly, we assessed the effect of the absence of accumulation in NE for reasoning-based seed inference with Seed2LP. Removing this constraint enabled to obtain seeds for all 107 GSMNs within the time limit, among which 106 harboured FBA-compliant solutions.

### Hybrid NE-FBA is a good trade-off between scalability and numerical accuracy for seed detection

We assessed the relevance of reasoning and hybrid solving modes of Seed2LP on the same benchmark of 107 GSMNs, with a time limit of 45 min or a limit of 1000 seed solutions computed. The use-case of this benchmark was to ensure the reachability of biomass reactants (reasoning and hybrid modes) and a positive flux in the biomass reaction (hybrid modes), objectives that are more biologically-relevant than reaching the whole set of metabolites. A naive implementation of a FBA mode imposing no NE reachability constraint and randomly selecting sets of seeds for satisfiability of linear constraints was also tested as a control.

Results are depicted in Table 1. Both *FBA naive* and *Hybrid-lpx* modes returned no seed solution before the timeout, indicating that these modes are not applicable to genome scale. The shared feature of *FBA naive* and *Hybrid-lpx* is the use of the hybrid ASP-simplex clingo-lpx solver: alternative hybrid modes in Seed2LP rather generate solutions with ASP and validate FBA directly during the solving (*Hybrid-GC)* or *a posteriori* (*Hybrid-filter*).

**Table 1.** Seed2LP results on 107 GSMNs (Target seed searching mode, subset minimal solutions) with several inference methods. The first column indicates the number of GSMNs for which solutions are obtained. The second (resp. third) column indicates the numbers of GSMNs for which at least one (resp. all) solution(s) enable the biomass reaction to carry flux in FBA.

| Solving mode | Nb. of net. with sol. | Nb. of net. with $\geq 1$ sol. FBA | Nb of net with all sol. FBA |
| --- | --- | --- | --- |
| *Reasoning* | 107 | 73 | 10 |
| *Hybrid-filter* | 71 | 71 | 71 |
| *Hybrid-GC* | 90 | 90 | 90 |
| *Hybrid-GC<sub>div</sub>* | 98 | 98 | 98 |
| *Hybrid-lpx* | 0 | 0 | 0 |
| *FBA naive* | 0 | 0 | 0 |

Several criteria can be used to evaluate the relevance of seed inference: obtaining one or several solutions that ensure the NE reachability of biomass reactants, ensuring a positive flux in the biomass reaction, or both of those. Results in Table 1 indicate that the *Reasoning* mode scales the best with solutions obtained for all GSMNs, and 1000 solutions for each. Hybrid modes found solutions for a smaller number of metabolic networks before timeout, between 66.4% (*Hybrid-filter*) and 91.6% (*Hybrid-GC*_*div*_).

All solutions provided by the hybrid modes succeed at FBA validation, as expected by design. *Hybrid-GC* method which adds constraints to the ASP solver to exclude previous sets of seeds (and their supersets) from future solutions helps to obtain more GSMNs with solutions (+19 wrt to *Hybrid-filter*). Increasing the diversity of solution contents with *Hybrid-GC*_*div*_ further increases the number of GSMNs having solutions (+8 wrt to *Hybrid-GC*).

The reasoning mode does not ensure FBA validation: only 9.3% of GSMNs exhibited a positive flux in all of their solutions. However for 68.2%, at least one solution out of up to 1000 had a positive flux, highlighting that this mode is not only scalable but can also provide FBA-compliant solutions. Identifying the latter requires an *a posteriori* validation with COBRApy though, which is implemented in Seed2LP. Including this validation in the solving, as performed by the *Hybrid-filter* increases the computational time, which explains the smaller number of GSMNs with a solution for the latter mode. We detail in Supplementary Material the computational time needed for the computation of a single solution in all four inference modes.

Above results were obtained with a subset-minimal optimisation. It is possible to use a minimise optimisation in Seed2LP but overall results on the benchmark led to less GSMNs having a solution, and less solutions by GSMN in 45 minutes. In this setting, *Hybrid-lpx* provided two solutions for the core network of *Escherichia coli* (95 reactions and 72 metabolites) using the minimise solving mode. Finally, we compare in Supplementary Material seed inference results obtained with Seed2LP or with an alternative flavour of the NE algorithm, ensuring self-regeneration of cycles in GSMNs.

### Exploration of seed diversity in solutions

We analysed further the content of seeds inferred by Seed2LP and the differences across inference modes by focusing on the GSMN of *Acinetobacter baumannii* AYE (iCN718) (888 metabolites, 1015 reactions). Again, the metabolic objective was the production of biomass precursors (reasoning and hybrid modes) in addition to a positive flux in the biomass reaction (hybrid modes). In the benchmark results (45 min or 1000 solutions) presented in Table 1, the reasoning mode raised 1000 solutions (465 FBA-validated), whereas *Hybrid-filter* (H_*F*_), *Hybrid-GC* (H_*GC*_) and *Hybrid-GC*_*div*_ (H_*GCdiv*_) only generated 149, 116 and 111 respectively before the timeout. While the sizes of seed solutions are relatively small (between 9 and 27 metabolites), enumerating solutions enables diverse metabolites to be selected as seeds, which can be biologically relevant for experimental validation. As such, the union of metabolites occurring in the solutions described above was of size 161, 67, 98, and 118 for reasoning, H_*F*_, H_*GC*_ and H_*GCdiv*_ respectively, highlighting that the latter mode indeed reaches more diverse solutions than the other hybrid modes.

In order to survey the diversity of seed solutions captured with larger enumerations, we ran Seed2LP until reaching 2000 solutions for each mode. It is worth noting that while reasoning computed this number of solutions in 6 seconds on a laptop, it took 8 days on a computing cluster for the *Hybrid-GC* mode to do so. However, only 381 solutions validated FBA among the reasoning ones. Sizes of the union of solutions show a low increase compared to the 45 min benchmark (199, 117, 135 and 149 for reasoning, H_*F*_, H_*GC*_ and H_*GCdiv*_ respectively), indicating that while the problem of seed inference has a high combinatorics, the number of metabolites involved in solutions is relatively small and does not increase much in larger enumerations. The relevance of *Hybrid-GC*_*div*_ over the other hybrid modes is observable in both the 45-minute and 2000-solution benchmarks: after the reasoning mode, it is the one that raises the largest union of solutions and therefore explores the most the solution space (Fig. 3a, Supplementary Material).

**Fig. 3.**
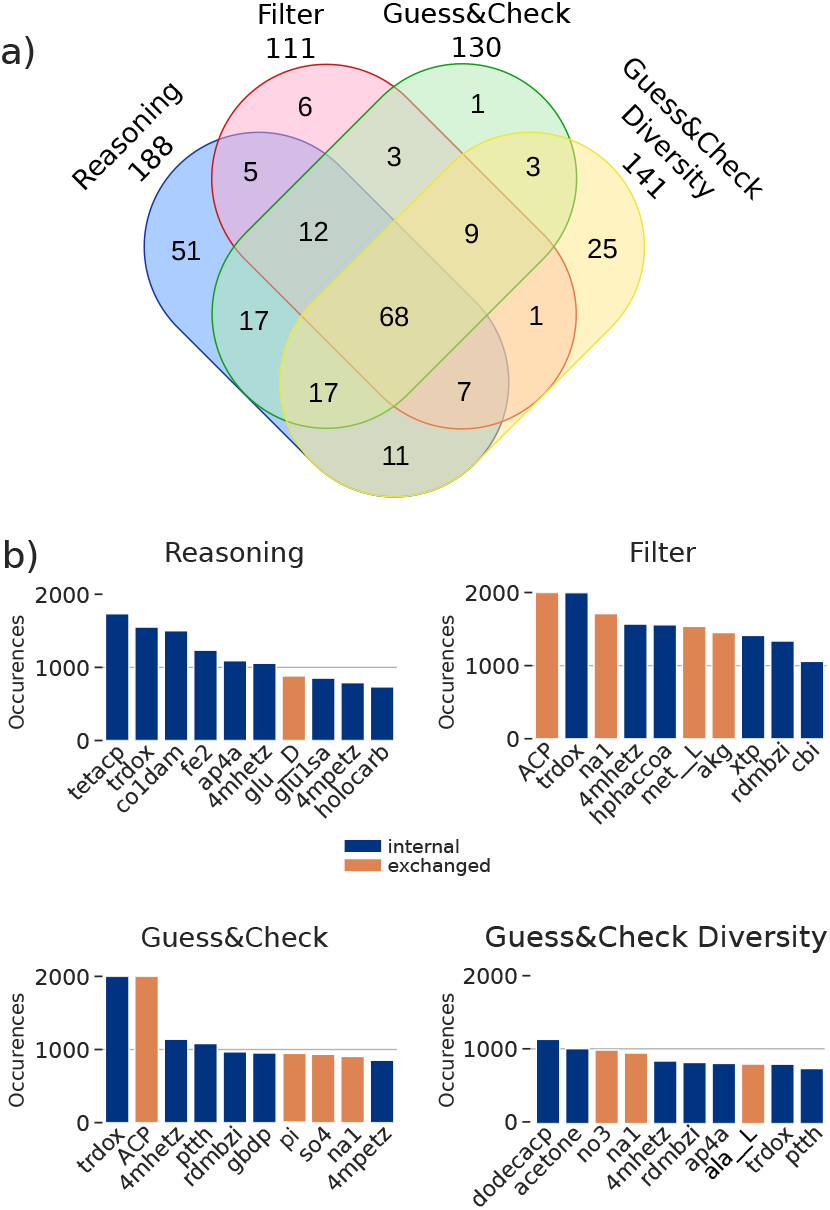
Seed inference in the iCN718 GSMN. **a)** Venn diagram illustrating the contents of solution unions in the four Seed2LP inference modes in 2000 solutions. **b)** Barplots depicting the 10 most frequent metabolites in seed solutions from the four modes. Orange metabolites correspond to molecules originally imported in the BiGG model, blue metabolites are internal. In both subfigures, seed unions were dereplicated before analysis, i.e., the same molecule in two different compartments will be considered identical.

We then surveyed the most frequent seeds across all solutions and the four modes, distinguishing internal metabolites from those which were imported in the original model (n = 120 metabolites imported through exchange or sink reactions). Figure 3b depicts the 10 most frequent metabolites per mode. Only one exchanged metabolite occurs among the most frequent one in reasoning mode whereas 3 or 4 appear in the hybrid modes. We can argue that the solutions validating the FBA propose more exchanged metabolites in the solutions than the *Reasoning*. The emphasis on diversity in the *Hybrid-GC* and especially *Hybrid-GC*_*div*_ modes is visible as few of the most frequent metabolites occur more than a thousand solutions (see also Supplementary Material, Supp. Fig. S3, Supp. Fig. S4). It is worth noting that the medium encoded in the original GSMN enables reaching 31% of metabolites with NE, and no biomass reactant, an observation that can be explained by the steady state modelling of FBA, in opposition to the more stringent initial-state modelling of NE [Frioux et al., 2019].

## Discussion

We developed a seed detection tool, Seed2LP, relying on a Boolean approximation model of metabolic producibility, that can be combined with FBA to infer growth medium nutrients. Obtaining chemically-defined or synthetic media has multiple applications in microbiology, especially for fastidious and uncultured microorganisms [Stewart, 2012]. Microbial dark matter, identified with techniques such as metagenomics impedes the understanding of most ecosystems harbouring a wide diversity of organisms [Mu et al., 2021, Kapinusova et al., 2023], but other use cases exist, for instance in the context of endosymbiont bacteria that cannot be grown outside of their hosts [Masson and Lemaitre, 2020]. Solving the culturability bottleneck has multiple applications, given the wide range of metabolic functions with biotechnological interest that microorganisms can provide [Rämä and Quandt, 2021, Lewis et al., 2021]. Systems biology and reverse ecology, through the implementation of dedicated computational models, can help addressing these objectives by suggesting necessary nutrients for growth.

Seed2LP is very flexible, allowing the user to choose between different seed search methods, focusing solely on the producibility of reactants of an objective reaction, or on the reachability of all metabolites in the network. Multiple parameters enable constraining seed inference, and solving modes based on reasoning or hybrid NE-FBA customise further the search. The reasoning NE-based model is highly scalable and efficient, often providing a thousand solutions well before the 45 min timeout of our benchmark, all ensuring that reactants of the objective reaction are in the scope. We chose the ability of models to carry flux in FBA as a validation method, an approach that is widely acknowledged in systems biology. We showed that this method outperforms other existing methods such as graph methods (NetSeed) or alternative NE flavours (Supplementary Material). Even if constraint-based modelling such as FBA is independent of network expansion, we observe that for the majority of GSMNs in both *Target* and *Full Network* modes, at least a solution ensures a positive flux in the biomass reaction. Hybridising NE with FBA ensures biomass production, but with a computational cost that leads less solutions to be generated, and a smaller solution space to be explored. We demonstrate nonetheless that a first solution can be obtained within a few seconds for most GSMNs. Reasoning and hybrid modes achieve varying level of diversity during seed enumeration, and among the hybrid one, *Hybrid-GC*_*div*_ reaches the most diverse solution contents.

The topology of GSMNs is very diverse and can facilitate or complicate the inference of seeds, resulting in solving modes that can be more or less efficient. As such, there is no optimal combination of parameters suitable for any GSMN, and one may benefit from testing and comparing several of them. While the inference can be fully automatised and performed systematically on a large collection of GSMNs, Seed2LP’s flexibility aims at making it useful in practice for experimental applications. It can be used in a back-and-forth process with bench work to partly automatise metabolic modelling-based medium inference, a use-case that typically requires curation [Nev et al., 2023, Tejera et al., 2020]. Selecting seeds among a subset of metabolites or forbidding some molecules to be used as seeds may prove useful in such iterative process. Seed2LP is a support to find the nutrient-associated missing link among the many factors of the environment that impact culturability: nutrients, pH, osmotic conditions, temperature, and others [Stewart, 2012]. A limitation of Seed2LP is therefore that it only considers metabolism to predict growth. Impact of molecule concentration, regulation processes or the dynamics of metabolism are additional facets whose analyses could be carried with more complex computational models, for instance in between seed inference and experimentation.

From a formal standpoint, our experiments suggest that the pure Boolean constraints of NE together with non-accumulation constraints form an efficient abstraction of the complete hybrid NE-FBA problem, with a limited false positive rate. Future work may investigate further discrete constraints that would make the over-approximation more accurate.

While use cases presented in this paper concern individual populations, the question of seed inference can be extended to co-culture contexts [Stewart, 2012], where interactions among species and division of labour can impact the minimal requirements of the community [Lewis et al., 2021]. Seed inference in that case could answer several objectives, ranging from stabilising or guiding the community to designing medium by taking cooperation into account or favouring it.

## Material and methods

### Benchmark Data

Curated GSMNs were retrieved from BiGG Database [Norsigian et al., 2019], using the API v2 last updated on 2019-10-31. The collection includes 108 GSMNs, but iAT PLT 636 was discarded from the analyses because of a lack of biomass reaction. GSMNs account mostly for bacteria, but also a few eukaryotes, including the human. All GSMNs have a set of exchange reactions that we can associate to seeds. 105/107 also hold a positive flux in FBA when optimising their biomass reaction, the two exceptions being Recon1 and iCN900.

### NetSeed

NetSeed analyses the topology of the graph associated to a GSMN in order to identify seeds. It computes the strongly connected components (SCCs) from the simple graph of the GSMN, constructs the directed acyclic graph (DAG) out of the SCCs and selects SCCs without input edges in the DAG. If those SCCs contain a single metabolite, it will be a seed, whereas if it contains several, one among them has to be selected as seed.

We downloaded the NetSeed-Perl tool from the NetSeed Website^2^ on 2022-01-19 [Carr and Borenstein, 2012]. GSMNs were normalised with Seed2LP (default parameters) prior NetSeed execution in order to ensure consistency between the network structure and associated mathematical model, and to remove exchange reactions. NetSeed was run with default parameters. It outputs a set of metabolites corresponding to the intersection of all solutions, sets of metabolites among which a seed must be selected. We performed a scalar product of the sets composing seeds to get the first thousand solutions.

### Analyses with Seed2LP

Seed2LP is a Python (v. 3.10) package, and that uses as main dependencies: Clyngor (v. 0.3.18) and Clingo-lpx (v. 1.3.0) for managing ASP, COBRApy (v. 0.26.0) [Ebrahim et al., 2013], MeneTools (v. 3.4.0) [Aite et al., 2018] and Metage2Metabo (v. 1.5.4) [Belcour et al., 2020].

Seed2LP was run with the 107 BiGG GSMNs, using the *full-network* and *target* objective modes, reasoning and hybrid inference modes. Unless stated otherwise, the subset-minimal optimisation was used and the reasoning modes used the *no-accumulation* setting. Unless stated otherwise, the execution of the programme was stopped when 1000 solutions were obtained, or after 45 min. An additional benchmark was performed with a 45-minute timeout to assess the time necessary to obtain one solution, with 1 core and 10Gb memory.

Validation of a positive flux in the biomass reaction was performed with COBRApy within Seed2LP for both Seed2LP and NetSeed solutions. The NE scope was computed through the Seed2LP *scope* functionality, using the SBML corresponding to the corrected GSMN. The presence of all GSMN metabolites (i.e., those occurring in reactions) in the scope was verified.

GSMN ICN718 was selected for further analysis and Seed2LP was run until 2000 solutions were obtained in *reasoning, Hybrid-filter* /*Hybrid-GC* /*Hybrid-GC*_*div*_ modes.

Calculation of exchanged metabolites coverage in inferred seeds consists in divising the number of exchanged metabolites (from exchange or sink reactions that import the molecule) among seeds by the total number of exchanged metabolites.

### Statistics and visualisation

All plots were generated with seaborn (v. 0.13.2) [Waskom, 2021, Hunter, 2007], using Python scripts to extract data, pandas (v. 2.2.2) [Wes McKinney, 2010] to aggregate data and manipulate tables and SciPy (v. 1.14.0) [Virtanen et al., 2020] for statistical tests. Unless stated otherwise, the latter were performed with the Kruskal-Wallis test [Kruskal and Wallis, 1952]. In all boxplots, the box indicates the quartiles of the dataset while the whiskers extend to the rest of the distribution, except for the points determined as outliers. The Venn diagramm was computed with http://bioinformatics.psb.ugent.be/webtools/Venn/ after retrieving the ID’s of the union of metabolites for each method stripped from their prefix and compartment suffix.

## Availability of code and data

Seed2LP is available on https://github.com/bioasp/seed2lp. Tools, datasets, analysis scripts and results are available on https://doi.org/10.57745/OS1JND.

## Author contributions statement

Conceptualisation: CF, LB - Supervision: CF, LP - Investigation: CF, CGN, LP - Formal Analysis: CF, CGN, LP - Software: CGN, CF, LB, LP, MB - Methodology: CF, CGN, LB, LP, MB - Validation: CF, CGN, LP - Visualisation: CF, CGN - Writing, Original Draft Preparation: CF, CGN, LP - Writing, Review & Editing: all - Funding acquisition: CF, LP.

## Acknowledgments

CF was supported by the French National Research Agency (ANR) France 2030 PEPR Agroécologie et Numérique MISTIC ANR-22-PEAE-0011. CF and LP were also supported by ANR in the scope of the project REBON (ANR-23-CE45-0008). Experiments presented in this paper were carried out using the PlaFRIM experimental testbed (https://www.plafrim.fr), supported by Inria, CNRS (LaBRI and IMB), Université de Bordeaux, Bordeaux INP and Conseil Régional de la Nouvelle Aquitaine. The authors acknowledge Sven Thiele for his valuable insights and helpful discussions.

A CC-BY public copyright license has been applied by the authors to the present document and will be applied to all subsequent versions up to the Author Accepted Manuscript arising from this submission, in accordance with the open access conditions of our funding.

## Supplementary Material

### Computational time for the inference of the first solution

Figure S1 illustrates the discrepancies in solving time across the four inference modes and in different *Full Network* or *Target* search modes. Results indicate that *Hybrid-filter* takes the most time to retrieve solutions, especially in *Target* search mode.

**Fig. S1.**
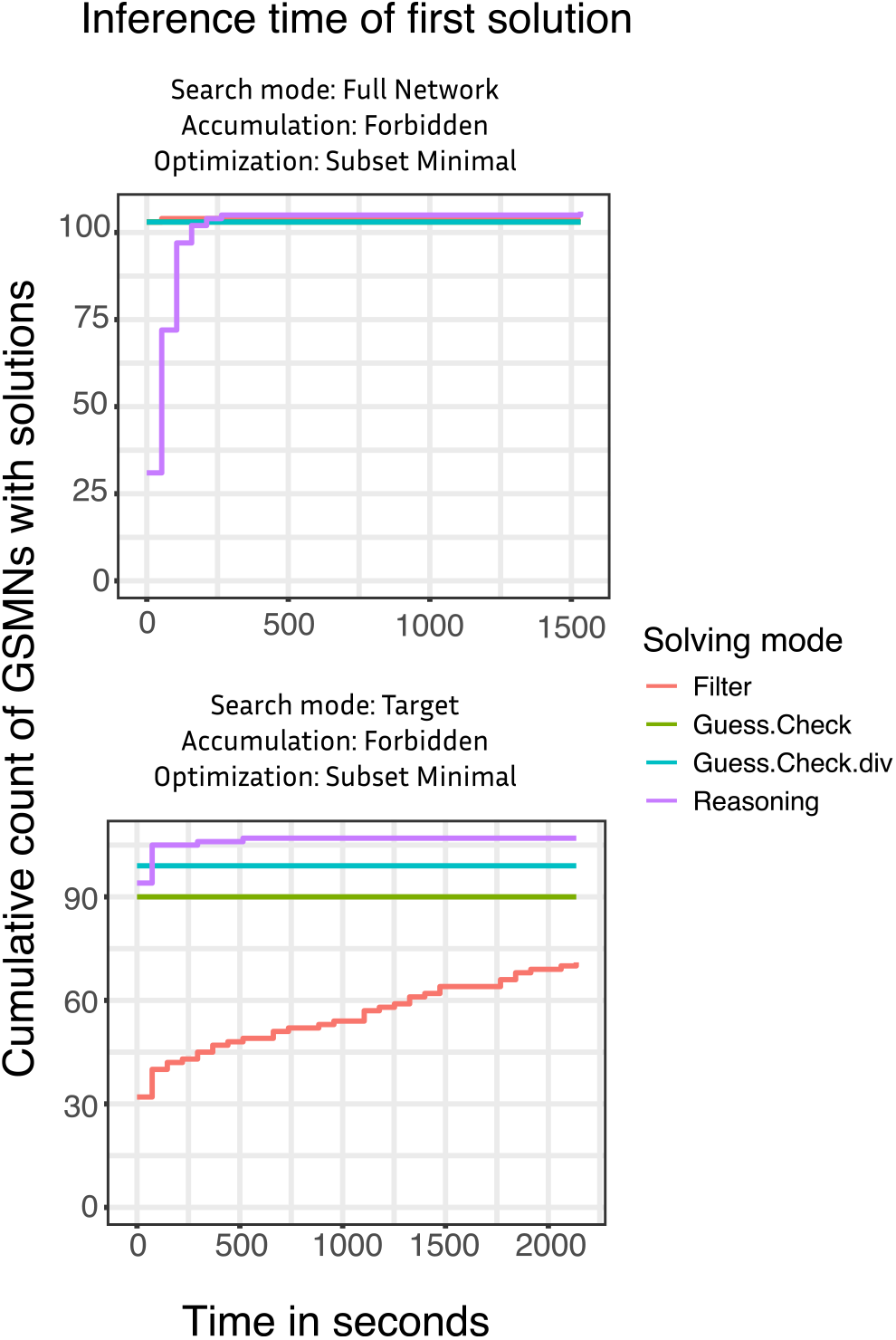
Solving time for the inference of a single seed solution in *reasoning, Hybrid-lpx, Hybrid-filter, Hybrid-GC* or *Hybrid-GC*_*div*_ modes. The cumulated number of GSMNs with a solution over time and before a 45 minute timout is illustrated. *Full Network* and *Target* modes are tested.

### Subset Minimal versus minimal-size optimisations

Finding minimal set of solutions is more computationally expensive than finding subset minimal solutions. We compared the two optimisations using all seed searching modes (*Full Network* or Target) and all seed inference methods (*Reasoning, Hybrid-lpx, Hybrid-filter, Hybrid-GC* or *Hybrid-GC*_*div*_). We analysed the sizes of seed solutions with respect to the size of the GSMN, and the coverage of exchange metabolites by sets of seeds. As the minimise optimisation does not raise solutions for all GSMNs, we restricted our analyses to GSMNs for which both optimisations gave results. We averaged all solution sizes per GSMN and compared the distribution of these values for the two optimisation modes with Kruskal-Wallis tests.

The proportion of exchange metabolites covered by seeds does not significantly change in subset-minimal versus minimal solutions regardless of the seed searching modes and inference method used. No significant difference occurs in the size of seed solutions in *Full Network* mode. In *Target* mode, only reasoning inference provides significantly smaller sets of seeds (n = 12, *χ*^2^ = 4.81, p = 0.028).

Overall, subset-minimal and minimal optimisations provide very similar results. The added-value in scalability associated with the subset minimal mode justifies its default usage in Seed2LP.

### Impact of NE flavours

We mentioned in the introduction that an alternative to NE takes into account self-regenerating cycles in an attempt to resemble, although qualitatively, the approach of FBA [Cottret et al., 2008, Acuña et al., 2012]. An ASP-based implementation of seed solving with such an underlying model is available with the Precursor tool (https://github.com/bioasp/precursor). We compared Seed2LP with Precursor in order to assess the impact of NE flavour on qualitative seed inference.

#### Methods

We ran Precursor (commit a3acf85) on the 107 GSMNs of the BiGG database with a 45 minutes time limit, either in *Target* mode, i.e., setting up all biomass reactants as targets, or *Full Network* mode, i.e., setting up all GSMN metabolites as targets. We additionally tested the impact of Seed2LP network-model reconciliation prior to the Precursor run.

#### Results

We first refer to Precursor results with *a priori* reconciliation provided by Seed2LP (Fig. S2). At least one solution was obtained before timeout for all GSMNs in *Full Network* mode, and 11 of them held a positive biomass flux in FBA. Interestingly, those latter were the same ones as those holding a positive flux in the NetSeed experiment (Fig. 2), suggesting shared topological characteristics among them. Precursor raised solutions for all GSMNs in *Target* mode as well, although no one was FBA compliant (Fig. S2a). An interesting observation is the relatively small size of Precursor’s seed sets with respect to the size of the network that arises from the consideration of self-regenerating cycles in GSMNs (Fig. S2b). Precursor generates sets of seeds ranging from 6 to 285 metabolites in *Full Network* mode, and 1 to 18 in *Target* mode whereas Seed2LP solutions contain between 6 and 59 seeds in *Target* mode and *Reasoning* method. Less solutions are found by Precursor than by Seed2LP: 82% of GSMNs have less than 8 solutions in 45 minutes in *Target* mode. Finally, we took a closer look at iCN718, as we analysed it further with Seed2LP. In *Full Network* mode, Precursor found a unique solution before timeout for this GSMN, consisting in 141 metabolites and enabling a positive flux in the biomass reaction. 246 solutions of size 1 (n = 245) or 2 (n = 1) seeds where found in *Target* mode, none satisfying FBA constraints. The impact of normalisation was null regarding the FBA results. It only slightly changed the number of solutions raised by Precursor for some GSMNs.

Overall, even if the network expansion flavour modelled in Seed2LP does not account for self-regenerating cycles, results from its reasoning modes (Fig. 2 for *Full Network* mode, Table 1 for *Target* mode) are more compatible with FBA than Precursor.

**Fig. S2.**
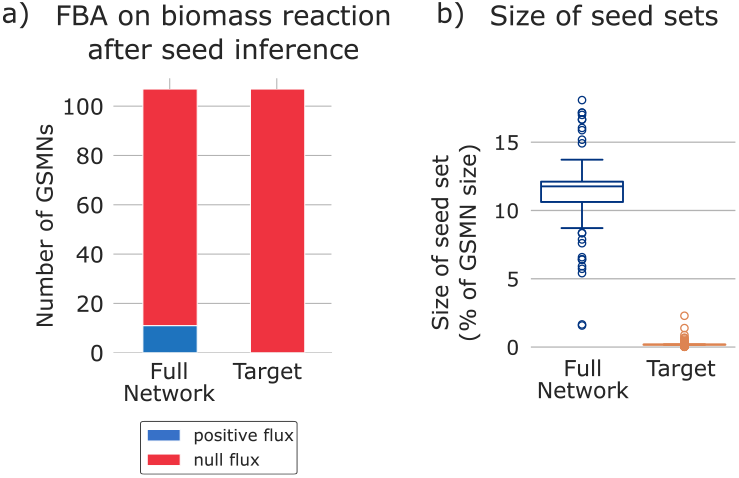
Results of Precursor seed inference. a) Number of GSMNs having all solutions validating FBA (blue) or none (red) or without any solution (black) in *Full Network* or *Target* search modes. b) Size of seed sets described as the proportion of the total number of metabolite per GSMN. Distribution of the average proportions computed by GSMN if several solutions were obtained for a given GSMN.

### Supplementary results on the iCN718 analysis

Table S1 provides solving details on the inference of seeds for the iCN718 GSMN, either in the 45-minute or 1000 solution set-up, or after 2000 solutions.

**Table S1.** Solving-associated details for seed inference in iCN718. First row depicts the results in the main benchmark presented in the manuscript (45 minute timeout or stop after 1000 computed solutions). Second row illustrates the results for computing 2000 solutions in each mode. The number of rejected models corresponds to solutions that were discarded because they did not validate FBA. Union and intersection describe the number of metabolites occurring in at least one solution and all solutions respectively. Abbreviations: R, *Reasoning*; F, *Hybrid-filter* ; GC, *Hybrid-GC* ; GCD, *Hybrid-GC*_*div*_; TO, Timeout; Nb, Number.

| Test | Nb solutions | Nb rejected | Time | Union | Intersection |
| --- | --- | --- | --- | --- | --- |
| 45-minutes TO<br>or 1000 solutions | R: 1,000 | R: NA | R: 6.209s | R: 161 | R: 1 |
|  | F: 149 | F: 799 | F: 45m | F: 67 | F: 2 |
|  | GC: 116 | GC: 801 | GC: 45m | GC: 98 | GC: 2 |
|  | GCD: 111 | GCD: 811 | GCD: 45m | GCD: 118 | GCD: 0 |
| 2000 solutions | 2,000 | R: NA | R: 6.403s | R: 199 | R: 0 |
|  |  | F: 9,964 | F: ≤8d | F: 117 | F: 1 |
|  |  | GC: 14,368 | GC: ≤8d | GC: 135 | GC: 2 |
|  |  | GCD: 9,273 | GCD: ≤8d | GCD: 149 | GCD: 0 |

Figure S3 present the characteristics of seed solutions in the four inference modes. We observe that the number of exchanged metabolites covered by reasoning solutions is lower than for hybrid ones (Fig. S3a). The seed set sizes remain low in all four modes (Fig. S3b). 73% of all GSMN metabolites are reachable with sets of seeds in average, but small variation occurs depending on the inference mode (Fig. S3c).

Figure S4 presents the frequency of occurrence of seed metabolites across the 2000 solutions computed by the 4 inference modes of Seed2LP. We observe that exchanges metabolites (n = 128) as described in the original SBML model are more frequently present in seeds computed with hybrid modes, suggesting that this characteristic may be associated to solutions which validate FBA. Nonetheless, as it grasps the most diverse set of metabolites, the reasoning mode is the one that retrieves the most exchanges metabolites within its union (37.5%).

**Fig. S3.**
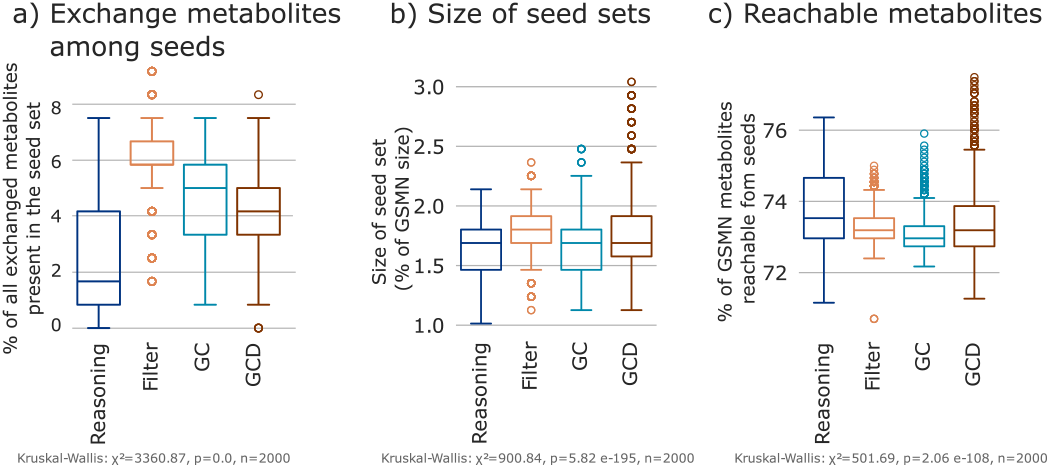
Characteristics of the 2000 solutions computed for iCN718: proportion of exchanged metabolites among seeds (a), size of seed sets with respect to the size of the GSMN (b), and proportion of reachable metabolites from the seeds (c), all according to the inference mode (*reasoning, Hybrid-lpx, Hybrid-filter, Hybrid-GC* or *Hybrid-GC*_*div*_).

**Fig. S4.**
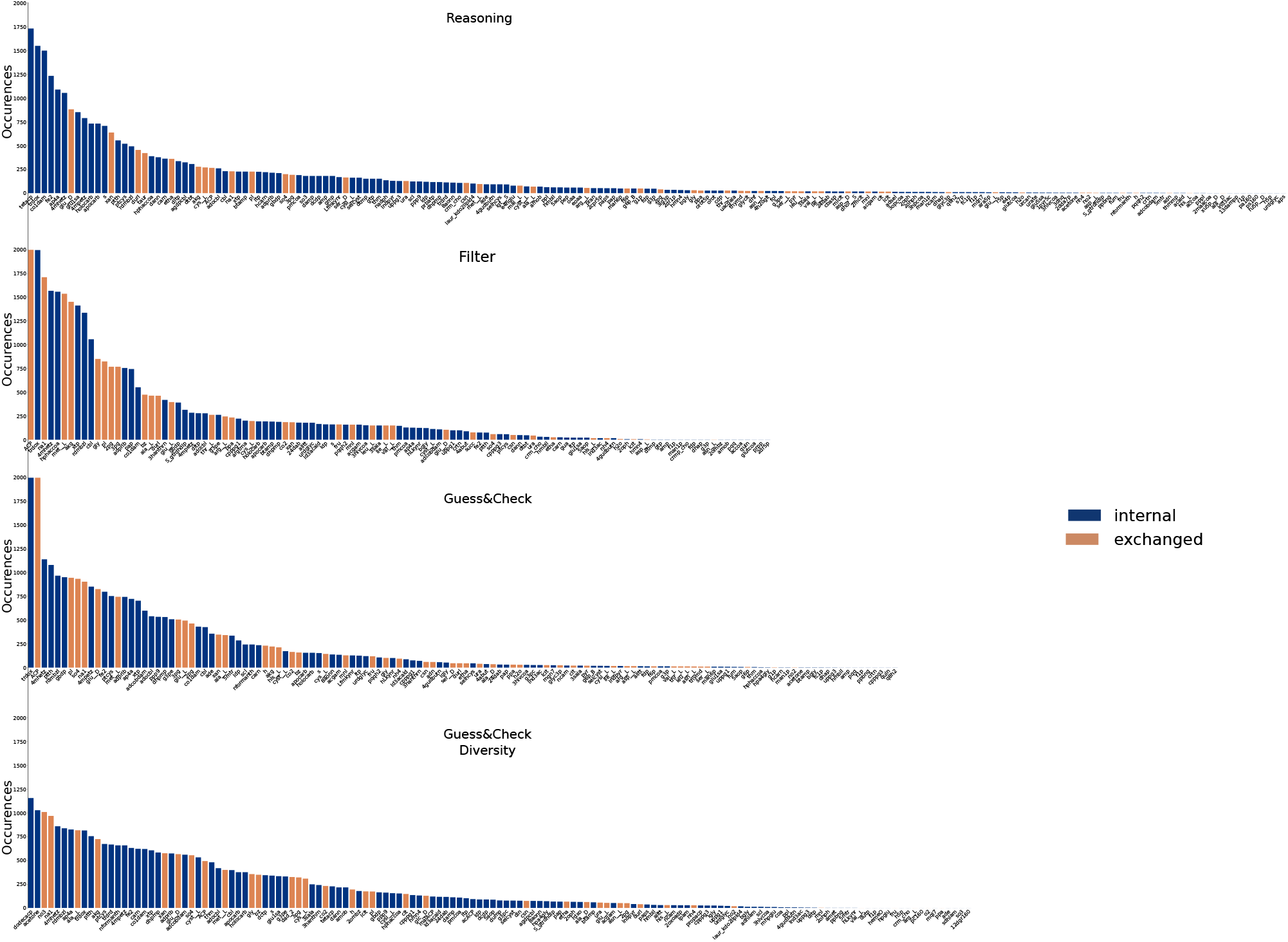
Occurrence of metabolites across the 2000 solutions according to the inference mode (*Reasoning, Hybrid-lpx, Hybrid-filter, Hybrid-GC* or *Hybrid-GC*_*div*_). Orange metabolites are imported (exchanged metabolites) in the original model, whereas blue ones are internal metabolites.

## Footnotes

1 https://github.com/potassco/clingo-lpx

2 http://borensteinlab.com/software_NetSeed.html

